# The human ventromedial prefrontal cortex: sulcal morphology and its influence on functional organization

**DOI:** 10.1101/417824

**Authors:** Alizée LOPEZ-PERSEM, Lennart Verhagen, Céline Amiez, Michael Petrides, Jérome Sallet

## Abstract

The ventromedial prefrontal cortex (vmPFC), which comprises several distinct cytoarchitectonic areas, is a key brain region supporting decision-making processes and it has been shown to be one of the main hubs of the Default Mode Network, a network classically activated during resting state. We here examined the inter-individual variability in the vmPFC sulcal morphology in 57 humans (37 females) and demonstrated that the presence/absence of the inferior rostral sulcus and the subgenual intralimbic sulcus influences significantly the sulcal organization of this region. Furthermore, the sulcal organization influences the location of the vmPFC peak of the Default Mode Network, demonstrating that the location of functional activity can be affected by local sulcal patterns. These results are critical for the investigation of the function of the vmPFC and show that taking into account the sulcal variability might be essential to guide the interpretation of neuroimaging studies.

**SIGNIFICANCE STATEMENT:** The ventromedial prefrontal cortex (vmPFC) is one of the main hubs of the Default Mode Network and plays a central role in value coding and decision-making. The present study provides a complete description of the inter-individual variability of anatomical morphology of this large portion of prefrontal cortex and its relation to functional organization. We have shown that two supplementary medial sulci predominantly determine the organization of the vmPFC, which in turn affect the location of the functional peak of activity in this region. Those results show that taking into account the variability in sulcal patterns might be essential to guide the interpretation of neuroimaging studies of the human brain and of the vmPFC in particular.

## INTRODUCTION

The ventromedial prefrontal cortex (vmPFC) is a key region for decision-making and subjective valuation processes (Lebreton et al., 2009; Noonan et al., 2010; Lopez-Persem et al., 2016). However, this brain region is not only involved in reward and value processing but also in emotional regulation (Hänsel and von Känel, 2008), memory representation (Bonnici et al., 2012) and it is a main hub of the Default Mode Network (Raichle, 2015), a network classically observed during resting-state and mind-wandering. Thus, vmPFC function is now the focus of many studies.

Defining the functional contributions of a cortical territory requires first an understanding of its anatomical organization. The discrete cytoarchitectonic areas found in a region have been shown to have distinct connectional profiles with other cortical and subcortical areas and also to relate to separate functional processes (Petrides and Baddeley, 1996; Petrides, 2002). These cytoarchitectonic areas often relate to sulcal patterns that can be clearly visualized in functional imaging studies. Indeed, the functional contributions of brain areas are constrained by their inputs and outputs, their connectivity patterns (Passingham et al., 2002; Petrides, 2002).

Characterizing the functions of the vmPFC is challenging for two reasons. First, the term ‘vmPFC’ does not refer to a specific anatomically delineated brain area. In functional neuroimaging studies that are based on the average brain activity of a group, the term ‘vmPFC’ has been used to label a large portion of the prefrontal cortex (Bartra et al., 2013) that includes several cytoarchitectonic areas, i.e. areas 10, 14, 25 and 32 (Figure 1) and vmPFC boundaries are debated (Wallis, 2011). The second challenge in assigning a function to the different subdivisions of the vmPFC comes from the fact that the classic neuroimaging approach is to average results across subjects in the Montreal Neurological Institute (MNI) stereotactic space (Evans et al., 1993). This approach ignores the considerable interindividual variability of the morphological sulcal patterns of the vmPFC (Chiavaras and Petrides, 2000; Mackey and Petrides, 2014). It should be emphasized here that there is evidence in the vmPFC that cytoarchitectonic areas relate well to particular sulci (Mackey and Petrides, 2014).

**Figure 1:**
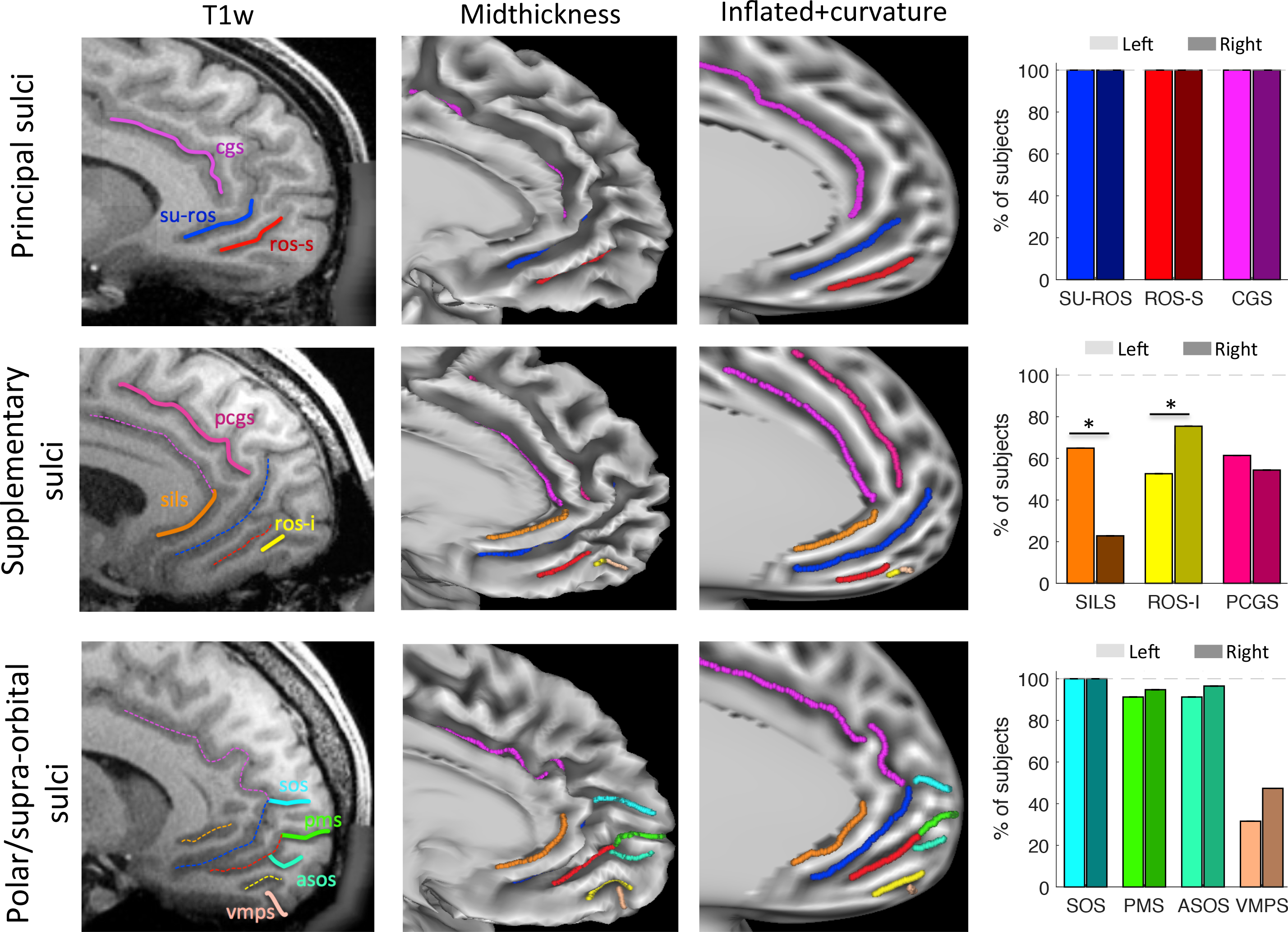
The ventromedial prefrontal cortex sulci. The sulci of the *ventromedial prefrontal cortex* (vmPFC) were identified using three image renderings: T1w volumes (first column), mid-thickness surface (2nd column) and inflated surface with the curvature value (3rd column). First row: the two principal sulci: the supra-rostral sulcus (SU-ROS) and the superior rostral sulcus (ROS-S). Second row: The inferior rostral sulci (ROS-I) and the subgenual intralimbic sulcus (SILS) are supplementary sulci. Third row: The supra-orbital sulcus (SOS), the accessory supra-orbital sulcus (ASOS), the polar medial sulcus (PMS) and the ventral medial polar sulcus (VMPS) composed the polar and supra-orbital sulci. Proportions of subjects exhibiting each sulcus are displayed in the fourth column for left (light colours) and right (dark colours) hemispheres. The cingulate sulcus (CGS) and the paracingulate sulcus (PCGS) are displayed for comparison purposes. Stars indicate significant differences between left and right hemispheres. On the volumes, solid lines represent the sulci of interest targeted in the fourth column. The dashed lines represent sulci identified in the preceding rows.

Sulcal patterns could indeed vary either in terms of presence/absence or in terms of shape / relative position of sulci. Importantly, it has been shown that the locations of distinct experienced value signals could be predicted from the organization of the sulci in the orbitofrontal cortex (Li et al., 2015). Moreover, taking into account the variability in sulcal patterns has proven to be essential to guide the interpretation of neuroimaging studies in the lateral prefrontal cortex and the cingulate cortex (Amiez et al., 2006, 2013). Thus, differences in functional properties in the vmPFC could also be related to the morphological heterogeneity across individuals.

In the present study, we first provide a precise description of the sulci of the vmPFC and we then show that distinct morphological patterns exist between individuals, preventing exact morphological alignment across individuals, and how those patterns affect the sulcal organization of the vmPFC. Finally, we show that this information is relevant to the study of the function of the vmPFC through a link between sulcal anatomy and the local response of the Default Mode Network.

## MATERIALS AND METHODS

### Subjects

The data used in this study are released as part of the Human Connectome Project (WU-Minn Consortium: Human Connectome Project, RRID:SCR_008749, http://db.humanconnectome.org) (Van Essen et al., 2012). We selected the S900 subjects release with 7T structural and resting-state fMRI data. The data were preprocessed according to the HCP pipeline (Glasser et al., 2013). All analyses were conducted on the data aligned using areal-feature-based registration (called “MSMAll” for “Multimodal Surface Matching” (Glasser et al., 2016)). This procedure aligns vertices on the cortical surface across subjects not only according to gross folding morphology, but also takes into account the subject-specific functional features, such as the location and distribution of resting-state networks. The MSMAll approach dramatically improves the functional alignment of cortical areas over and above registration based solely on volumetric or surface-based morphological registration. This type of registration is referred to as “area-based” registration and is sometimes considered a near optimal functional alignment (Glasser et al., 2016). Here we test whether considering detailed inter-individual variability in sulcal patterns can explain functional organisation even beyond this state-of-the-art area-based alignment approach. Of the 73 subjects of this specific HCP release, 16 subjects were excluded because of family ties with other subjects of the database. The data analysis was therefore based on 114 hemispheres of 57 subjects (37 females).

### Sulcal identification

The analyses were restricted to the ventral part of the medial wall of the prefrontal cortex, delineated by an arbitrary horizontal line that runs from the front of the brain to the genu of the corpus callosum. This delineation allowed us to define a broad anatomical mask used in the resting-state analysis. Following Mackey and Petrides, 2014, we identified the two main sulci in this region: the suprarostral sulcus (SU-ROS) and the superior rostral sulcus (ROS-S). An additional sulcus, ventral to the ROS-S, was observed in several subjects, and was labelled the inferior rostral sulcus (ROS-I).

Fronto-polar sulci, identified as MPS (medial polar sulci) in Mackey and Petrides, 2014, were dissociated into three potential sulci: ASOS (accessory supra-orbital sulcus), MPS (medial polar sulci) and SOS (supra-orbital sulcus) (see also <u>Petrides, 2018)</u>. We distinguished them as follows: the MPS is formed by a fold in the fronto-polar cortex creating a sulcus on the medial wall while ASOS and SOS were defined as folds on the medial wall itself. SOS was more dorsal than ASOS and PMS. An additional Ventral Medial Polar Sulcus (VMPS) was identified by the fact that it is the most ventral and directed towards the orbital part of the cortex, dorsal to the olfactory sulcus.

Although the cingulate cortex was not the focus of the present study, the cingulate sulci were also identified because they are necessary to define the morphological patterns of the ventromedial sulci. The main cingulate sulcus (CGS) was identified in all subjects, but the para-cingulate sulcus (PCGS), a secondary sulcus, was sometimes observed either as a well-marked sulcus or as a trace of it, that we labelled as spur-PCGS. Finally, we identify the SILS (subgenual intralimbic sulcus). This sulcus lies below the genu, hence the term “subgenual”, and is located between the SU-ROS and the corpus callosum.

The cortical surface renderings were generated using the Connectome Workbench viewer (RRID:SCR_008750, http://www.humanconnectome.org/connectome/connectome-workbench.html) (Marcus et al., 2011). In order to label each sulcus observed in the region of interest in every hemisphere, we examined three types of renderings of the medial wall: the volume (T1w), the mid-thickness surface and the dilated surface with a projection of the curvature (Figure 1). We then manually drew the sulcus on the dilated surface following the curvature projection.

### Morphological analysis

All analyses and statistics were conducted in Matlab 2017a (MATLAB and Statistics Toolbox Release 2017a, The MathWorks, Inc., Natick, Massachusetts, United States, RRID:SCR_001622, http://www.mathworks.com/products/matlab/) with in-house bespoke scripts calling Workbench executables. To compute the statistics and get information on each type of sulcus, regions of interest were created from the manual drawing explained above for every hemisphere. MNI coordinates (Montreal Neurological Institute, bic-mni-models, RRID:SCR_014087) were accessible in all the surface files provided in the HCP dataset. For display purposes, in Figure 4E, we used a dilated version of the sulci (5mm larger) for the analysis on the normalized length in order to increase the number of points entered in the analysis. The results did not change when the original sulcus was used as a region of interest (posterior versus anterior part weight comparison: t(113)=−3.98, p=1.10^−4^).

To compute the probability of observing one sulcus on one side given its presence on the other side, we used the following equation:

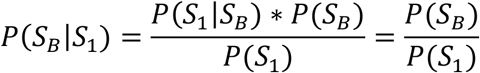

With P(S_B_) the probability of sulcus on both sides, P(S1) the probability of sulcus on only one side. P(S_1_|S_B_)=1.

If not specified in the text, all comparisons were assessed by two-tailed t-tests. Analysis on volumes were conducted using FSL (RRID:SCR_002823).

### Resting state analysis

The rs-fMRI acquisitions (including the use of leading-edge, customized MRI hardware and acquisition software) and image processing are covered in detail in <u>Glasser et al., 2013; Smith et al., 2013; Uğurbil et al., 2013</u>. After image preprocessing (primarily using the FMRIB Software Library, FSL, RRID:SCR_002823) (Jenkinson et al., 2012), FreeSurfer (RRID:SCR_001847)(Fischl, 2012), and Connectome Workbench (Marcus et al., 2013, software packages), the functional timeseries are filtered and artefacts are removed using an automated data-driven approach that relies on ICA decomposition and hand-trained hierarchical classification (FMRIB’s ICA-based X-noisifier [FIX]) (Smith et al., 2013).

After these image preprocessing and timeseries cleaning steps, the resting-state fMRI data enter a probabilistic independent component analysis (MELODIC, <u>Beckmann and Smith, 2004)</u> to identify statistically independent spatial component maps and their accompanying timeseries. These spatial maps are not binary and can be overlapping: the value at each point in the brain can be considered a ‘weight’ reflecting how strongly this point contributes to a given component. As a whole, each signal component map can be considered as a statistically independent probabilistic parcel reflecting a specific brain network. As such, the component weight maps are not the same as a seed-based correlation map but are often interpreted in analogous ways.

To identify the component capturing the Default Mode Network, we examined them one by one for every subject and, for each session, we extracted the component that had the highest weights in the precuneus and posterior cingulate cortex, in bilateral posterior parietal cortex and in bilateral temporal association cortex, following the definition provided by <u>Raichle (2015)</u>. Weights correspond to the probability of each vertex belonging to a specific IC. Note that in order to avoid any bias in our results, we ignored the weights on the medial wall of the prefrontal cortex (both the vmPFC and the dorso-medial prefrontal cortex). When several components were matching those criteria, we selected the one that was most bilateral. Components identified as DMN were finally averaged across sessions to obtain one DMN per subject.

### Definition of the regions of interest (ROI)

In the last section of the results, we compare four ways of selecting a region of interest in the vmPFC.

In the first method, called ‘a priori’, the ROI was defined from a meta-analysis done on <u>Neurosynth (www.neurosynth.org, RRID:SCR_006798)</u> with the term ‘Default Mode Network’ as a search term. The MNI coordinates of the peak of the Default Mode Network in the vmPFC were used to define the region of interest on each subject as a disc of diameter 10mm around it. This ROI is an MNI-based ROI: vertices on MSMAll surfaces are different across subjects but the MNI coordinates are the same.

The second and the third methods are labelled ‘Circular inference’ methods because they are selected on the basis of the group result and then tested (they will provide positive results by definition). The ‘group’ ROI was defined according to the averaged DMN weight map across the group of subjects. We set an arbitrary threshold of weights to obtain a clear cluster in the vmPFC and then select this cluster as an ROI. This ROI is a vertex-based ROI (MNI-coordinates are different across subjects).

The ‘peaks’ ROI was based on the same map as the previous ROI but instead of selecting a cluster, the 10 vertices with the highest DMN weights were selected. MNI coordinates of those vertices were extracted and back-projected on each subject to extract the DMN weights at the subject level. This ROI is an MNI-based ROI.

Finally, the anatomical ROI, called anterior ROS-S (aROS-S), was determined at the subject level based on the ROS-S drawn in the previous analysis. For each subject, we extracted the DMN weights within the anterior half of the ROS-S. This ROI is an anatomy-based ROI (MNI and vertices will be different across subjects).

For every subject, the DMN weights were extracted within each ROI and averaged across the ROI in order to get one value per ROI and per subject. Then, the significance of the differences between ROIs at the group level was assessed with paired t-tests.

In order to compare the quality of DMN identified with the ‘a priori’ method and with the anatomical method, we defined three regions of interest representative of the DMN: the Posterior Cingulate Cortex (PCC), the anterior Superior Temporal Sulcus (aSTS) and the Temporo-Parietal Junction (TPJ). We drew them manually on the map of averaged DMN weights (based on a threshold of 20 arbitrary units of weight).

Signal consistency corresponds to (1/STD_ROI), with STD_ROI corresponding to the standard deviation of the time series across vertices within each ROI. A low standard deviation indicates small differences of the signal across vertices of an ROI. We represented 1/STD_ROI in figure 7.

## RESULTS

### Description of the sulci

Using a subset of 57 subjects (i.e. 114 hemispheres) from the Human Connectome Project (see Methods), we identified all the possible sulci within the left and right hemispheres of the ventral part of the medial wall of the Prefrontal Cortex, i.e. the ventro-medial Prefrontal Cortex (vmPFC). Three categories of sulci were observed: the principal medial sulci, the supplementary medial sulci and the polar/supra-orbital sulci (Figure 1 and Table 1).

**Table 1.**
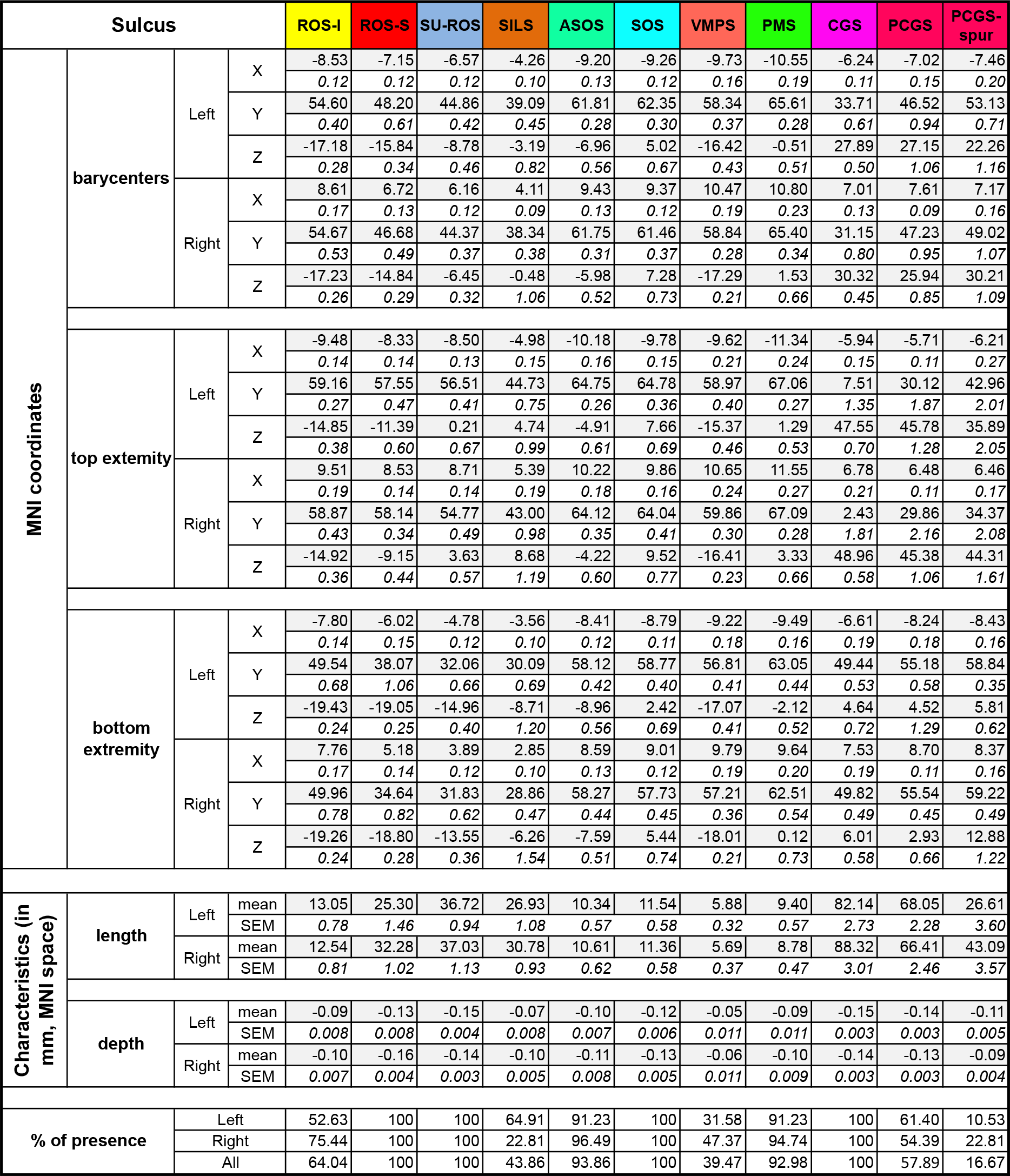
Summary description of the vmPFC sulci. Average (grey highlighted) and SEM (italic) MNI coordinates of the barycentre, top extremity and bottom extremity of each vmPFC sulcus on each hemisphere across subjects. Length, depth and percentage of presence are also reported. ‘Top’ corresponds to the location of the most antero-dorsal (postero-dorsal for cingulate sulci) extremity of each sulcus, ‘Bottom’ corresponds to the location of the most ventro-posterior (antero-ventral for cingulate sulci) extremity of each sulcus. SU-ROS: Suprarostral sulcus; ROS-S: superior rostral sulcus; SILS: subgenual intralimbic sulcus; ROS-I: inferior rostral sulcus; SOS: supra-orbital sulcus, ASOS: accessory supra-orbital sulcus; PMS: polar medial sulcus; VMPS: ventro-polar sulcus; CGS: cingulate sulcus; PCGS: paracingulate sulcus; PCGS-spur: spur paracingulate sulcus

#### Principal medial sulci

The supra-rostral sulcus (SU-ROS) and the superior rostral sulcus (ROS-S) were identified in all hemispheres. The mean average length of the SU-ROS was 36.87±0.73 mm with no significant difference between the left and right hemispheres (left: 36.71±0.94 mm, right: 37.03±1.13 mm; left vs right: t(112)=−0.21, p=0.83, unpaired t-test). The ROS-S had an average length of 28.78±0.95 mm with a significant difference between the left and right hemispheres (left: ROS-S is 25.29±1.46 mm, right: 32.27±1.02 mm; left vs right: t(112)=−3.92, p=2.10^−4^, unpaired t-test). We did not find any significant correlation between the length of the sulci on the two hemispheres across subjects (SU-ROS: r=−0.13, p=0.35, ROS-S: r=0.20, p=0.13).

At the group level, we observed an overlap in the MNI locations of the centre of gravity of those two sulci. The ROS-S average location was y=48.20(±0.61), z=−15.84 (±0.34) on the left side and y=46.68 (±0.49), z=−14.84 (±0.29) on the right side, and the SU-ROS average location was y=44.86 (±0.42), z=−8.78 (±0.46) on the left side and y=44.37 (±0.37), z=−6.45 (±0.32) on the right side. However, on the dorso-ventral axis, the range of location of the ROS-S on the left side varied from z=−21 to −9 mm and from z=−16 to 4 mm for the SU-ROS, indicating that, at the group level, there is an overlap in the MNI coordinates of these two sulci, notably between z=−16 and −9 mm. This result was also observed in the right hemisphere with a ROS-S range from z=−20 to −7 mm and a SU-ROS range from z=−12 to 0 mm.

#### Supplementary medial sulci

The inferior rostral sulcus (ROS-I) was present in 64.04% of the hemispheres, with 52.63% and 75.44% in the left and right hemispheres, respectively. The ROS-I was significantly more frequent in the right hemisphere compared with the left hemisphere (t(112)=−2.59, p=0.01, unpaired t-test). The ROS-I average location was y=54.60(±0.40), z=−17.18(±0.28) on the left side and y=54.67(±0.53), z=−17.23(±0.26) on the right side. The probability of having an ROS-I on both sides given its presence on one side or the other was estimated at 52.08%, suggesting a relative independence of the two hemispheres regarding this sulcus.

The length of the ROS-I was on average 12.74±0.70 mm with no difference between the two hemispheres (left: 13.04±1.07 mm; right: 12.54±0.93 mm; left vs right: t(71)=0.35, p=0.72, unpaired t-test). There was no correlation of length between the two hemispheres when it was present on both sides (r=0.11, p=0.59).

The subgenual intralimbic sulcus (SILS) was identified in 43.86% of the hemispheres, with 64.91% and 22.81% of the left and right hemispheres, respectively. The difference between hemispheres was significant (t(112)=4.96, p=3.10^−6^, unpaired t-test), i.e. it was more probable to find this sulcus in only one hemisphere compared to the two hemispheres (the probability that SILS is on both sides given its presence on one side was 19.05%). The average location of SILS was y=39.09(±0.45), z=−3.19(±0.82) on the left side and y=38.34(±0.38), z=−0.48(±1.06) on the right side. With an average length of 27.93±1.13 mm, there was no significant difference of the SILS length between the two hemispheres (t(48)=−1.51, p=0.14) and no correlation of length for the cases in which SILS was found in both hemisphere (r=0.27, p=0.52)

#### Polar and supra-orbital sulci

The supra-orbital sulcus (SOS) was found in all hemispheres. The accessory supra-orbital sulcus (ASOS) was present in 93.86% of the hemispheres with no over-representation in the right hemisphere: 96.49% in the right and 91.23% in the left hemisphere (t(112)=−1.17, p=0.25, unpaired t-test). Its existence on one side was predictive at 91.07% of its presence on both sides. The SOS average location was y=62.35(±0.30), z=5.02(±0.67) on the left side and y=61.46(±0.37), z=7.28(±0.73) on the right side. The ASOS average location was y=61.81(±0.28), z=−6.96(±0.56) on the left side and y=61.75(±0.31), z=−5.98(±0.52) on the right side.

The polar medial sulcus (PMS), found in 92.98% of the hemispheres, was not more frequent in one hemisphere compared with the other (left: 91.23%; right: 94.74%, t(112)=−0.73, p=0.47, unpaired t-test). Its existence on one side was predicted by its existence on the other side (85.96%). The PMS average location was y=65.61(±0.28), z=− 0.51(±0.51) on the left side and y=65.40(±0.34), z=1.53(±0.66) on the right side.

Finally, we identified in some subjects a small sickle-shaped sulcus just dorsal to the olfactory sulcus in the anterior ventral polar region. This ventral medial polar sulcus (VMPS) was observed in 39.47% of the hemispheres with a trend for a difference between the two hemispheres (left: 31.58%, right: 49,12%, left vs right: t(112)=−1.73, p=0.09, unpaired t-test). The probability of having this sulcus in both hemispheres given its presence on one of the hemispheres was 36.36%, suggesting that it is common to have this sulcus in only one hemisphere. The VMPS average centre of gravity was y=58.34(±0.37), z=−16.42(±0.43) on the left side and y=58.84(±0.28), z=−17.29(±0.21) on the right side.

### Description of patterns

We next examined the interactions between the different sulci. In agreement with Mackey and Petrides (2014), several interaction patterns between the SU-ROS and other sulci were identified. A fourth kind was also noted but it represented less than 5% of the subjects. We subdivided each group based on the presence or absence of a PCGS.

Type 1 corresponds to a merging of the SU-ROS with the cingulate sulcus (sub-type 1a) or with the paracingulate sulcus (sub-type 1b). In type 2, the SU-ROS merges with one or more polar sulci but not with the cingulate or paracingulate sulci. Subtypes were defined on the basis of the polar sulcus with which the SU-ROS merges (2a: SOS; 2b: ASOS; 2c: PMS; 2d: merges with more than one polar sulcus). Type 3 refers to a merging of the SU-ROS with both the cingulate sulcus and a polar/supra-orbital sulcus. Similar to subtypes identified in type 2, subtype 3a merges with the cingulate sulcus while subtype 3b merges with the paracingulate sulcus. These subtypes could also be separated in groups according to which polar sulcus the SU-ROS merges with (3a1: SOS, 3a2: ASOS, 3a3: PMS, 3a4: two polar sulci) (Figure 2A). Type 4 corresponds to a merging with both cingulate and paracingulate sulci (subtype 4a). It could also merge with polar sulci (subtype 4b) (Figure 2B).

**Figure 2:**
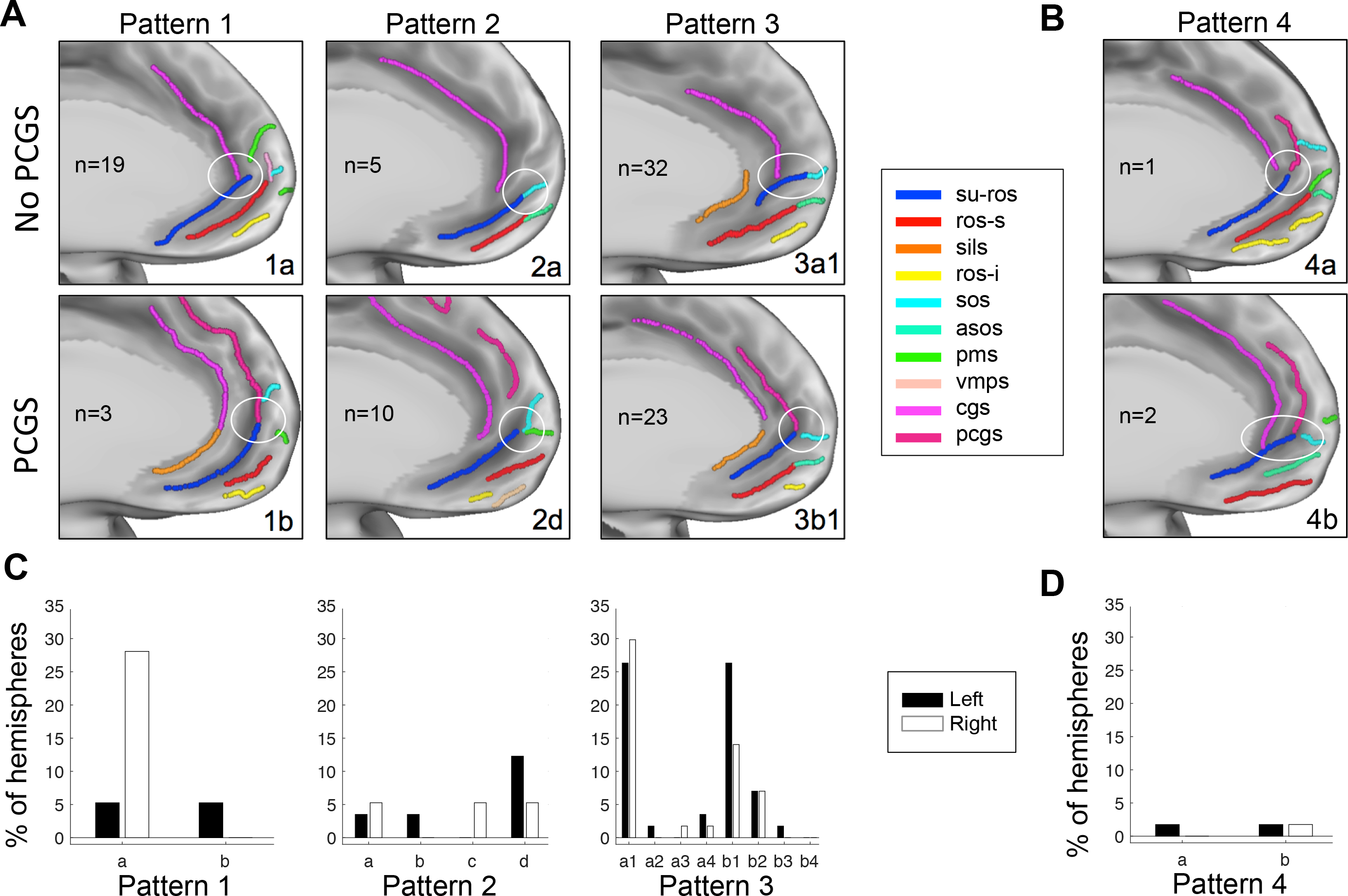
vmPFC anatomical patterns. **A.** Examples of the three common types of sulcal patterns in the vmPFC. The patterns depend on how the SU-ROS merges with other sulci (white circles) and the absence (first row) or absence (second row) of a paracingulate sulcus PCGS. In type 1, the SU-ROS merges with either the CGS (1a) or the PCGS (1b). In type 2, the SU-ROS merges with one or more polar/supra-orbital sulci but not with the CGS (2a) or the PCGS (2d). In type 3, the SU-ROS merges with both polar/supra-orbital sulcus and CGS (3a1) or PCGS (3b1). **B.** In the rare type 4, the SU-ROS merges with both CGS and PCGS, without (4a) or with (4b) a merging with a polar/supra-orbital sulcus. n indicates the number of hemispheres with each pattern. **C** and **D**. Percentage of left (black) and right (white) hemispheres displaying each pattern and sub-type of patterns. 1a: only CGS (left: 5.3%, right: 28.1%), 1b: only PCGS (left: 5.3%, right: 0%); 2a: only SOS(left: %, right: 5.3%), 2b: only ASOS (left: 3.5%, right: 0%), 2c: only PMS (left: 0%, right: 5.3%), 2d: two polar/supra-orbital sulci (left: 12.3%, right: 5.3%); 3a: with CGS and with SOS (3a1, left: 26.3%, right: 29.8%), or ASOS (3a2, left: 1.8%, right: 0%), or PMS (3a3, left: 0%, right: 1.8%), or with two polar/supra-orbital sulci (3a4, left: 3.5%, right: 1.8%). 3b: with PCGS and with SOS (3b1, left: 26.3%, right: 14.0%), or ASOS (3b2, left: 7.0%, right: 7.0%), or PMS (3b3, left: 1.8%, right: 0%) or two polar/supra-orbital sulci (3b4, left: 0%, right: 0%). 4a: CGS and PCGS (left: 1.8%, right: 0%); 4b: CGS, PCGS and polar/supra-orbital sulcus (left: 1.8%, right: 1.8%). SU-ROS: Suprarostral sulcus; ROS-S: superior rostral sulcus; SILS: subgenual intralimbic sulcus; ROS-I: inferior rostral sulcus; SOS: supra-orbital sulcus, ASOS: accessory supra-orbital sulcus; PMS: polar medial sulcus; VMPS: ventral medial polar sulcus; CGS: cingulate sulcus; PCGS: paracingulate sulcus.

In agreement with Mackey and Petrides (2014), we found that 19.3% of all hemispheres belong to type 1, 17.5% to type 2, and 60.5% to type 3. Type 4 was the smallest group with only 2.6% of hemispheres showing this pattern (Figure 2C & D).

### Morphological deformation associated with sulcal pattern variability

On the dorso-ventral axis, we observed that the barycentres of the ROS-S and SU-ROS in the left hemisphere spread over 11 and 8mm, respectively (13 and 12mm in the right hemisphere). In order to capture better how the inter-individual variability in sulcal patterns impacts the organization of the vmPFC, we extracted for each subject the antero-posterior and dorso-ventral MNI coordinates of the barycentre, the antero-dorsal top extremity, the postero-ventral bottom extremity and the length of the ROS-S and the SU-ROS. We computed the partial correlation coefficients between the 14 variables (Z and Y MNI for barycentre, top extremity, bottom extremity and sulcal length for each sulcus) and the following 6 factors: absence or presence of the SILS, ROS-I and PCGS and the assignment to pattern 1, 2 or 3. Significant influences were observed that were related to the presence of the SILS and ROS-I but not to the presence of the PCGS (all p>0.05). An effect of patterns 2 and 3 was also observed but nothing for pattern 1 (all p>0.05) (Figure 3A). We describe the significant results in the following part.

**Figure 3:**
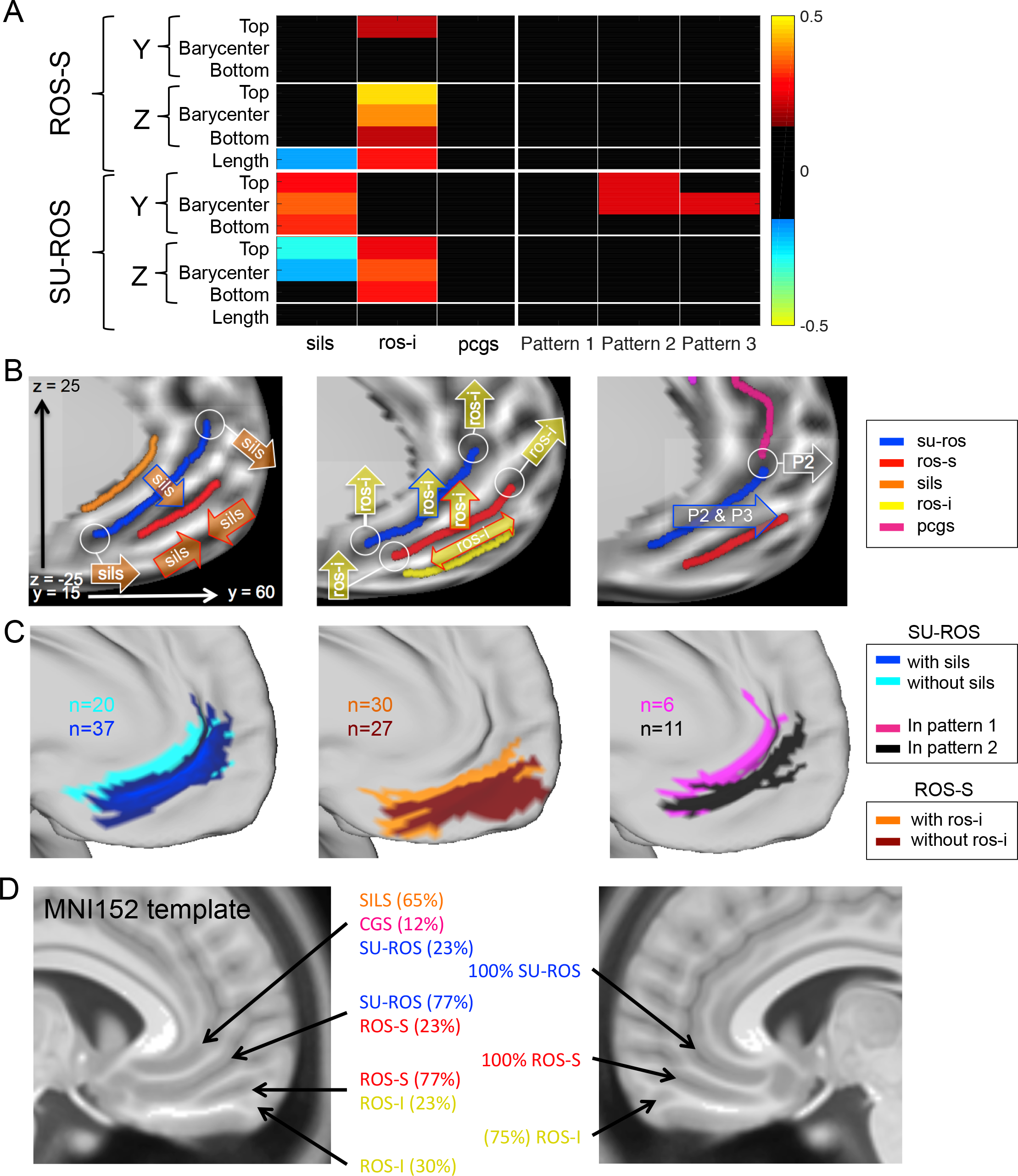
Morphological deformation associated with the presence of supplementary sulci and patterns. **A.** Partial correlation coefficients between each measure of interest of the ROS-S and SU-ROS and the presence of SILS, ROS-I, PCGS and patterns 1 to 3. Y and Z correspond to the antero-posterior and dorso-ventral MNI coordinates, respectively. ‘Top’ corresponds to the location of the most antero-dorsal extremity of each sulcus, ‘Bottom’ corresponds to the location of the most ventro-posterior extremity of each sulcus. ‘Length’ is the length of each sulcus. Only significant partial correlation coefficients are displayed (p<0.05). Hot colours indicate a positive influence of having a supplementary sulcus or a specific pattern. Cold colours indicate negative effects. **B.** Schematic representation of the significant influence of the presence of SILS (left), the presence of ROS-I (middle) and patterns 2 and 3 (P2 and P3, right) on the SU-ROS and ROS-S locations. Arrows attached to white circles indicate an influence on the circled extremity of the sulci; arrows with coloured bordures indicating an influence on the barycentres of the sulci; double arrows indicate a positive effect on length while two facing arrows indicate a negative effect on length. **C**. Left: Projection of all the left SU-ROS for subjects with and without SILS. Middle: Projection of all the left ROS-S for subjects with and without ROS-I. Right: Projection of all the left SU-ROS for subjects with pattern 1 and subjects with pattern 2. n indicates the number of subjects displayed on the surface. SU-ROS: Suprarostral sulcus; ROS-S: superior rostral sulcus; SILS: subgenual intralimbic Sulcus; ROS-I: inferior rostral sulcus; PCGS: paracingulate Sulcus. **C.** vmPFC sulci in the MNI152 template. On the left hemisphere, the sulci observed on the MNI152 template correspond to an overlap of several sulci. Percentages were computed from our sample of subjects. This overlap is induced by the morphological deformation caused by supplementary sulci. The right hemisphere of the MNI152 template does not present such overlaps because of the high presence of ROS-I. This absence of overlap does not mean that supplementary sulci do not induce morphological deformation on the right side; it indicates that the average template has sulci in the right hemisphere that globally correspond to the same right hemisphere sulci across subjects.

#### SILS

The presence of the SILS had a strong influence of the MNI location of the SU-ROS because it induced a global mean shift of 3.1mm (MNI-space) of the SU-ROS toward the antero-ventral direction. Indeed, the MNI Y and Z coordinates of the SU-ROS were shifted for the barycentre (Y: shift=2.5mm, rho=0.35, p=2.10^−4^; Z: shift=1.8mm, rho=−0.21, p=0.03) and the top extremity (Y: rho=0.28, p=3.10^−3^; Z: rho=−0.29, p=2.10^−3^). The bottom extremity was also shifted towards the anterior direction (Y: rho=0.30, p=1.10^−3^). Additionally, the ROS-S length was shorter when SILS was present (rho=−0.19, p=0.04) (Figure 3B).

#### ROS-I

The presence of ROS-I had a strong impact on the MNI location of both principal sulci: the presence of ROS-I was dorsally shifting the barycentre of the ROS-S (Z: shift=2.2mm, rho=0.39, p=3.10^−5^) and the SU-ROS (Z: shift=2.7, rho=0.34, p=3.10^−4^). This dorsal shift could also be seen in a drift of the bottom and top extremities of both sulci (Top SU-ROS: rho=0.24, p=0.01, ROS-S: rho=0.46, p=4.10^−7^; Bottom SU-ROS: rho=0.30, p=2.10^−3^, ROS-S: rho=0.21, p=0.03). An additional effect of the presence of ROS-I was found on the ROS-S length: it was longer in hemispheres with ROS-I compared to hemispheres without it (rho=0.29, p=2.10^−3^). This was also observed in a significant effect on the anterior location of the top extremity (rho=0.20, p=0.03) (Figure 3C).

As the presence or absence of ROS-I and SILS had opposite effects, we checked to what extent a hemisphere with ROS-I but without SILS would have a shift in its ROS-S and SU-ROS locations compared to a hemisphere without ROS-I but with SILS. The average antero-ventral shift in MNI space was 3.5mm and 4.2mm for the ROS-S and the SU-ROS, respectively. The maximal dorso-ventral distance observed between such two opposite hemispheres was 16.3mm (9.3 mm for the antero-posterior direction). This result shows that the deformations associated with sulcal morphology can be critical for the interpretation of activity based solely on MNI coordinates compared to an anatomical referential.

#### Patterns

We did not observe a significant effect of the PCGS on the ROS-S or the SU-ROS morphology. We therefore merged the different subgroups of patterns based on the presence or absence of a PCGS to test for an effect of patterns on ROS-S and SU-ROS morphology. We only observed an effect on the SU-ROS antero-posterior location. Indeed, the SU-ROS in pattern 2 was more anterior (Barycentre: shift=2.4mm, rho=0.24, p=0.01; Top extremity: rho=0.23, p=0.02). The pattern 3 also induced an anterior shift of the barycentre of the SU-ROS (shift=2.0mm, rho=0.23, p=0.02) (figure 3D).

To characterize further the feature that had the greatest influence on the sulcal morphological variability in the vmPFC, we carried out a Principal Component Analysis (PCA) on the 14 variables collected for every hemisphere. The first three components explained 84.1% of the variance in the data (47.4%, 24.8% and 11.9% for the first, second and third components). We extracted those three first components and ran again the partial correlation analysis against the six features of interest. The results demonstrated that the ROS-I was the only feature significantly affecting the first component (rho=0.31, p=9.10^−4^) and the SILS was the only feature affecting significantly the second component (rho=−0.27, p=5.10^−3^). The third component was significantly affected by the ROS-I presence (rho=0.29, p=2.10^−3^) but not by any other features (despite trends for pattern 2 and pattern 3, with rho=0.18, p=0.058 and rho=0.17, p=0.077 respectively). This result suggests that the ROS-I presence or absence can explain more than 59.3% of the variability in the data compared with at least 24.8% for the SILS presence.

As a control analysis, we checked whether hemispheres with and without ROS-I had undergone different deformation during the normalization procedure of the brain volumes. We computed the deformation (curvature extracted from T1w undistorted images versus curvature extracted from T1w but resampled on MSMAll normalized) for each hemisphere and then compared it between hemispheres with and without ROS-I. No significant difference was observed, suggesting that the MSMAll normalization procedure did not affect differentially subjects with and without ROS-I. Consequently, morphological deformation induced by ROS-I presence cannot be attributed to higher artefactual deformation induced by volume normalization.

Finally, in order to illustrate the impact of our result, we investigated how the principal and supplementary sulci were represented in the MNI152 template (Fonov et al., 2009), conventionally adopted in FSL (Jenkinson et al., 2012) and SPM (Friston, 2007). We registered original T1w volumes of our group of subjects on the MNI template (non-linear registration) and grouped subjects according to the presence or absence of ROS-I and SILS and observed that the sulci observed on the right hemispheres were corresponding to SU-ROS and ROS-S. However, on the left hemisphere of the template, the sulci do not correspond to a unique sulcus but to a mixture of sulci across individuals. Indeed, the first subgenual sulcus corresponds to an overlap of the SILS, the CGS and the SU-ROS, the sulcus below this one corresponds to an overlap of the SU-ROS and the ROS-S, and the more ventral sulcus corresponds to an overlap between the ROS-S and the ROS-I (Figure 3D). This phenomenon is primarily driven by the presence of ROS-I and SILS, which are close to 50% in the left hemisphere. Therefore, interpreting the location of activity in the left vmPFC regarding the sulci of the MNI template might be misleading and careful conclusions should be drawn from them.

### Peak of default mode network related to sulcus location

Finally, we tested whether the interindividual variability of the vmPFC sulcal pattern could impact its functional organization. This analysis was based on resting-state functional MRI and more specifically on a network associated with the vmPFC: the default mode network (DMN). The DMN is a network mainly composed of the vmPFC, the posterior cingulate cortex and the lateral parietal cortex (Raichle, 2015). It can be easily identified during resting state MRI (rs-fMRI) by applying Independent Component Analysis (ICA). In the HCP dataset used in the present study, for each subject, the ICA component corresponding to the DMN was identified (see methods). We then obtained for each subject a brain map of the DMN component. Each vertex of this map is associated with a weight, which corresponds to the probability of each vertex belonging to the DMN (Beckmann and Smith, 2004). An average of those weights is displayed in Figure 4A.

**Figure 4:**
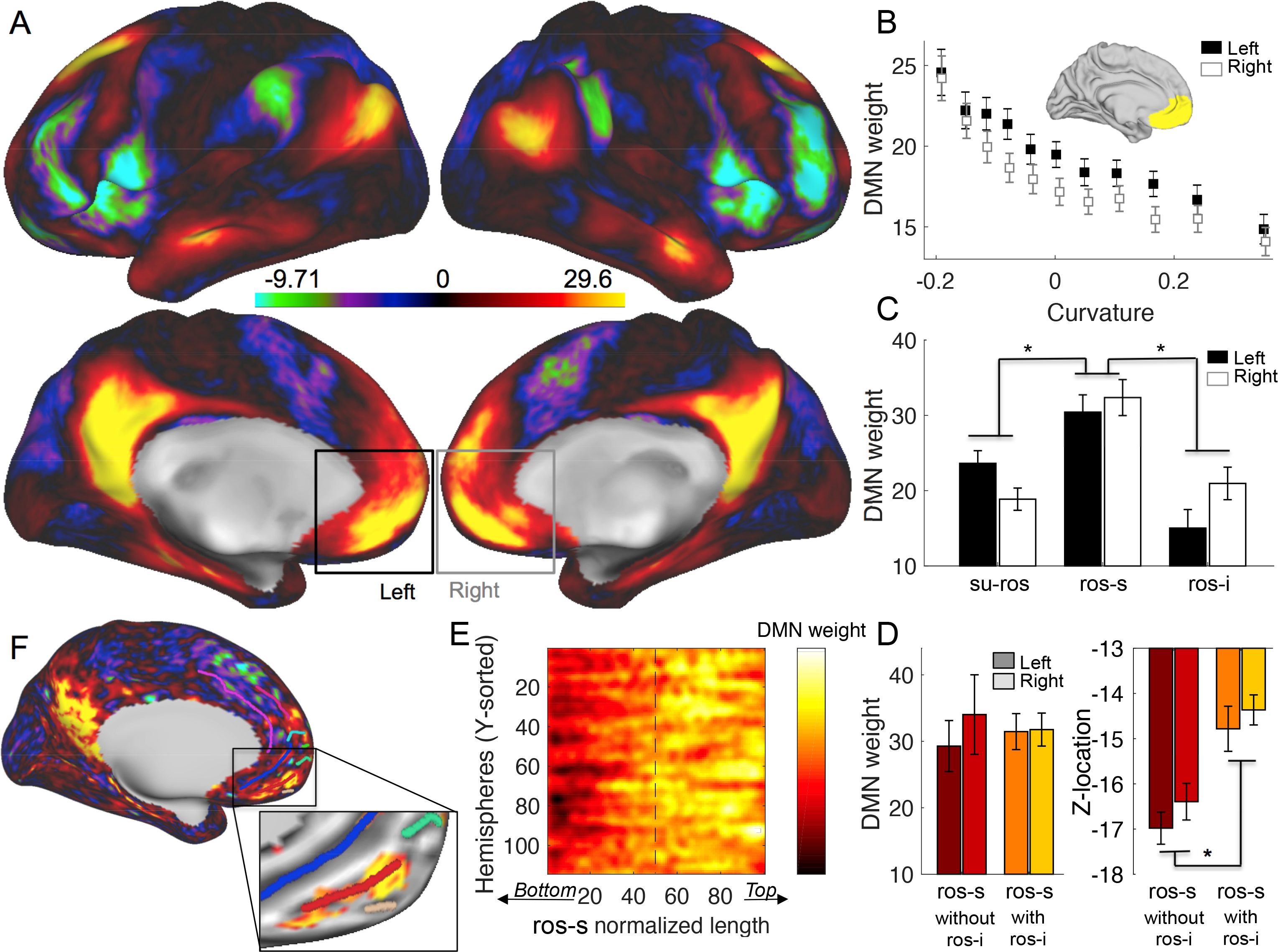
Default Mode Network weights and their dependence on the superior rostral sulcus. **A.** Default Mode Network weights averaged across subjects on each hemisphere. **B.** DMN weights are negatively correlated with the cortical curvature within the vmPFC (anatomical mask shown in the insert) for left (black) and right (white) hemispheres (squares represent binned data across hemispheres). Positive curvature corresponds to gyri while negative curvature corresponds to sulci. **C.** Averaged DMN weights across subjects in the SU-ROS, the ROS-S and the ROS-I in left (black) and right (white) sides. **D.** Averaged DMN weights (left) and Z-MNI location (right) in the ROS-S across subjects without ROS-I (red) and with ROS-I (orange) for left (dark colour) and right (light colour). Error bars represent SEM. **E.** Matrix of all ROS-S aligned on their normalized length and sorted according to their Y-MNI location. Hot colours indicate high DMN weights. **F.** Subject example of DMN with sulci. Insert is a zoom of the vmPFC with an arbitrary threshold of the DMN weights. The strongest DMN weights are located on the anterior part of the ROS-S (red). SU-ROS: Suprarostral sulcus; ROS-S: superior rostral sulcus; ROS-I: inferior rostral sulcus.

First, we determined whether vmPFC sulci-based ROIs would be relevant for the rs-fMRI analysis. We extracted the curvature (depth of the sulcus and gyrus) within a broad vmPFC mask (see methods). We regressed the DMN weights against the curvature within the vmPFC mask and found a strong negative effect across subjects (left: M=−19.1, SD=2.12, t(56)=−8.10, p=5.10^−11^; right: M=−20.5, SD=1.96, t(56)=−8.54, p=1.10^−11^)(Figure 4B), indicating that the DMN weights were higher in deep areas, i.e., in sulci. Critically, this result indicates that the sulci within the vmPFC contribute more to the Default Mode Network than the gyri. In other words, a lack of correlation between a vmPFC sulcus-based ROI and DMN weights cannot be attributed to a poor sulcal MRI signal as opposed the one recorded on the gyrus.

We then extracted the DMN weights from the principal medial sulci (SU-ROS and ROS-S) and the ROS-I and tested for a difference between them. We found an effect of sulcus in a one-way ANOVA (F(2,297)=18.7, p=2.10^−8^). Pairwise comparisons indicated that the ROS-S had significantly higher weights than the SU-ROS (t(226)=5.1, p=8.10^−7^, unpaired t-test) and than the ROS-I (t(184)=4.9, p=2.10^−6^, unpaired t-test). No effect of side was observed (all p>0.05). Thus, at the group level, the ROS-S was identified as the main hub of the vmPFC for the Default Mode Network (Figure 4C).

Given that the ROS-I had a strong influence on the location of the ROS-S, we checked the validity of the previous result with a complementary ANOVA analysis including two factors: ‘group’ and ‘sulcus’. The group factor was defined as subjects with and without ROS-I (1/0) and sulcus was defined as ROS-S and ROS-I (1/2). Again, we found a strong effect of sulcus (F(1,183)=20.11, p=1.10^−5^) but no effect of group (F(1,183)=0.04, p=0.85) (Figure 4D). This finding confirms the previous result showing that the ROS-S is a critical hub of the DMN independently of the existence of the secondary ROS-I sulcus.

Finally, we extracted the DMN weights within the ROS-S and sorted each vertex according to the Y-axis. Given that each ROS-S across hemispheres had a different length, we normalized the length of the sorted vector to fit a 100 points vector. We then merged all the normalized vectors in a single matrix and observed that the highest DMN weights were located in the anterior part of the ROS-S, starting around 50% of the ROS-S length (Figure 4E). A simple comparison between the first half of the sulcus and the second half confirmed this observation (t(113)=−8.13, p=6.10^−13^, paired t-test). A representative hemisphere of this result is depicted in Figure 4F. Thus, we identified that the ROS-S, and its anterior part in particular, is the hub of the vmPFC subcomponent of the DMN.

Given that ROS-I has a strong influence on the location of the ROS-S, one prediction following this result is that the DMN connectivity between subjects with and without ROS-I would be slightly different, especially in the vmPFC region. Indeed, if the vmPFC hub of the DMN is in the ROS-S, subjects with a ROS-I should have a stronger connectivity in the dorsal vmPFC. We averaged the DMN weights of subjects with and without ROS-I separately and then contrasted those maps (Figure 5). We could observe that this prediction was verified in the vmPFC in which we found significant clusters (larger than 20 mm^2^ and with T-values higher than 2.7) in both left and right hemispheres. Moreover, a cluster also appeared in the PCC, suggesting that morphological variability in the vmPFC might also induce functional variability in distant regions such as the PCC.

**Figure 5:**
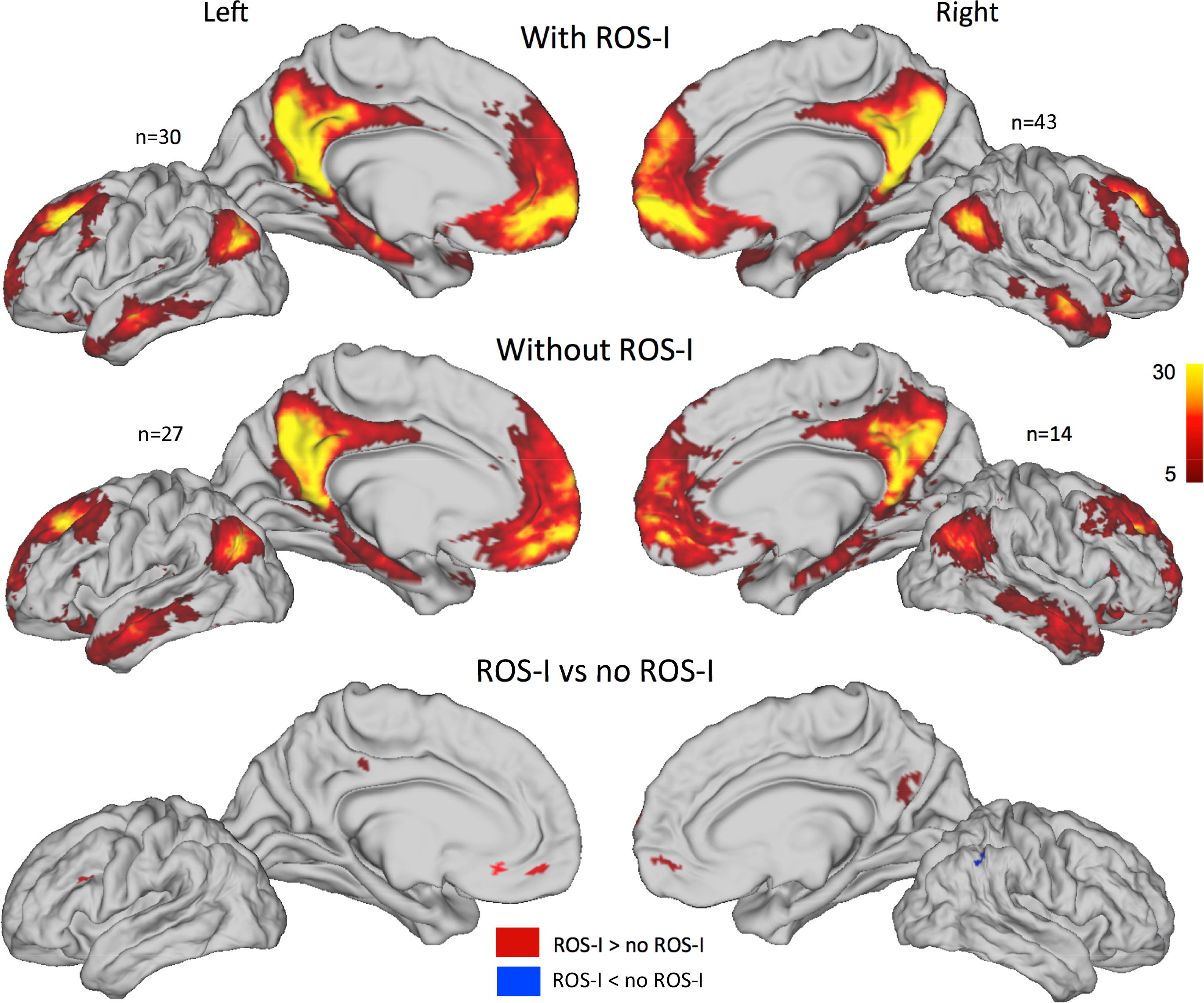
Default Mode Network weights in hemispheres with and without ROS-I. Default Mode Network (positive) weights averaged across subjects on each hemisphere for hemispheres with (top) and without (middle) ROS-I. n indicates the number of hemispheres in each averaged map. The contrasts of the two groups for each hemisphere are displayed in the bottom panel. Red (blue) clusters indicate higher (lower) weights for hemispheres with ROS-I compared to hemispheres without ROS-I. Clusters with t-value>2.7 and t-value<−2.7 and larger than 20 mm2 are depicted.

### Comparison of ROI definition methods to investigate the DMN

As we intend here to show that taking into account morphological features such as sulcal location might help the investigation of specific functions of the vmPFC, we tested whether using a region of interest defined according to sulcal morphology was better than using a sphere around specific MNI coordinates. We compared four ways of selecting a region of interest (ROI). The first one is an *a priori* ROI defined according to the peak of activity corresponding to the ‘Default Mode Network’ term search in neurosynth.com (y=50, z=−10): we labelled it ‘Prior’. The second and the third ones were defined from the average map of the Default Mode Network on the dataset used in our study and are depicted in Figure 4A. The second one corresponds to the cluster found on the average maps (Group cluster) and the third one corresponds to the 10 vertices with the highest DMN weights in the group (Group peaks). This procedure would be classically labelled as ‘double dipping’ or ‘circular inference’ since it is a selection of region following a positive result. It would bias the result in favour of a positive result. The fourth one is an anatomically based ROI: for each subject, we extracted the DMN weights of the anterior part of the ROS-S (aROS-S). A more detailed description of the ROI definition and weights extraction is included in the methods. All four ROIs had significant DMN weights (all p>0.05). Not surprisingly, the ‘Group’ ROI(s) showed a significantly higher effect than the ‘Prior’ ROI (Prior versus Cluster: t(113)=−6.43, p=3.10^− 9^, Prior versus Peaks: t(113)=4.52, p=2.10^−6^, paired t-tests). Interestingly we found that the anatomically defined ROI aROS-S had a significantly higher effect than the three other ROIs (paired t-tests against the ROS-S ROI: Prior: t(113)=6.72, p=8.10^−10^; Cluster: t(113)=2.74, p=7.10^−4^; Peaks: t(113)=3.57, p=5.10^−4^) (Figure 6). This result shows that using the anterior part of the ROS-S as a region of interest is more efficient at the group level than using MNI coordinates based on previous studies or on the group result. Thus, even when positively biasing an ROI selection, we have demonstrated that using anatomical information to select the ROI provides stronger results than using MNI coordinates as a reference.

**Figure 6:**
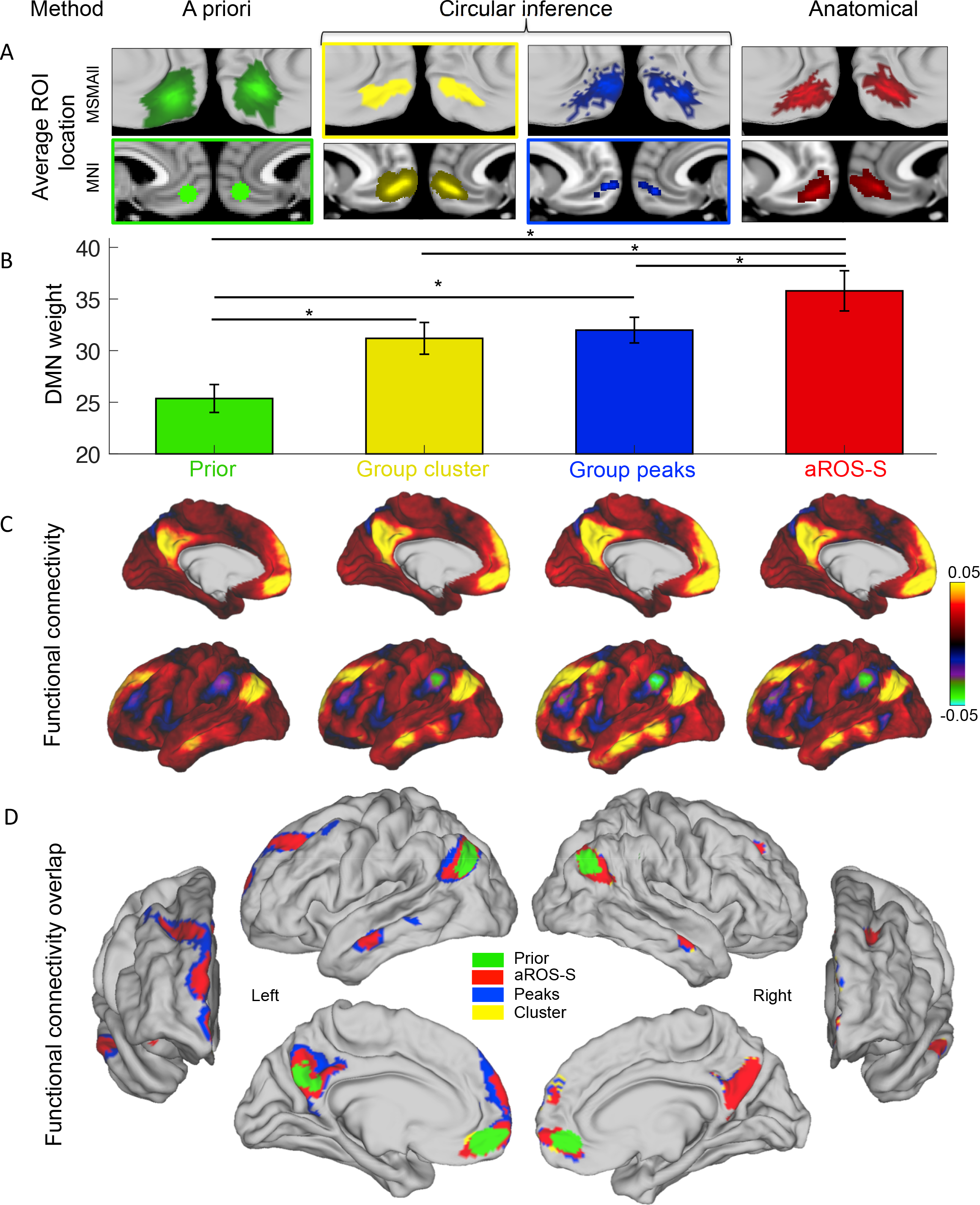
Default Mode Network weights in four vmPFC ROIs. DMN results from ROIs defined according to 4 different methods: a priori MNI method (green, ‘Prior’), circular inference method, based on group results (cluster, yellow and peaks, blue), and anatomical method (anterior ROS-S, aROS-S, red). **A.** Average ROI location across subjects on the MSMAll surfaces (top row) and in the MNI referential (bottom row). Coloured rectangles indicate the referential of each ROI: MNI for ‘Prior’ and ‘Group peaks’, vertices for ‘Group cluster’. There is no rectangle for the aROS-S because its referential is anatomical. **B.** Default Mode Network weights averaged across all hemispheres for each ROI. Error bars represent SEM. Stars indicate significance. **C.** Mean functional connectivity map across left hemispheres for each ROI. Colours indicate average correlation coefficients. Right hemispheres have a similar profile and are not displayed for illustrative purposes. **D.** Overlap of the maps displayed in C but thresholded at 0.05. Maps were superimposed according to the extent of the cluster (smaller above). The maps are displayed for left and right hemispheres with anterior, lateral and medial perspectives.

Finally, we evaluated the extent to which selecting the aROS-S as a region of interest was advantageous. First, we used the four previously described vmPFC ROIs as seeds to compute the functional connectivity of each region with the rest of the brain. We observed a very similar pattern of connectivity for all of them. However, when those maps are thresholded at 0.05 and overlapped, we can observe differences. The DMN observed with the Prior ROI is the smallest and the right PCC does not survive the thresholding. The ‘peaks’ and ‘cluster’ ROIs shared a very similar large network while the DMN observed with the aROS-S ROI was more restricted even when covering the same brain region. Thus, the DMN elicited by the aROS-S ROI has the best specificity among the four ROIs, allowing a clear separation of functional areas with the strongest evidence.

Then, we quantified the advantage of using aROS-S as a vmPFC ROI compared to an a priori ROI for the investigation of functional networks (Figure 7). We computed the average strength of connectivity of aROS-S and ‘Prior’ with the main other components of the DMN (TPJ, PCC and aSTS, see methods for ROI definition) and compared them. We found that the functional connectivity of the aROS-S with the three components was higher than the connectivity of the ‘Prior’ ROI (PCC: t(113)=6.62, p=1.10^−9^; TPJ: t(113)= 7.76, p=4.10^−12^; aSTS: t(113)= 7.02, p=2.10^−10^, paired t-tests). We then assessed the consistency of the recorded signal in the vmPFC ROI across the different vertices of each ROI. For each time point of the time series extracted from the aROS-S and the ‘Prior’, we computed the standard deviation across all vertices forming each ROI and then averaged it across time and hemispheres. The ‘signal consistency’, corresponding to the inverse of this measure was much stronger in the aROS-S (M=12.2; SD=0.11) compared to the ‘Prior’ ROI (M=6.7; SD=0.07) (t(113)=−20.66, p=4.10^−40^, paired t-test). Thus, by using the aROS-S as an anatomical ROI, we can have access to a more consistent signal in the vmPFC, but also to a well and clearly defined DMN.

**Figure 7:**
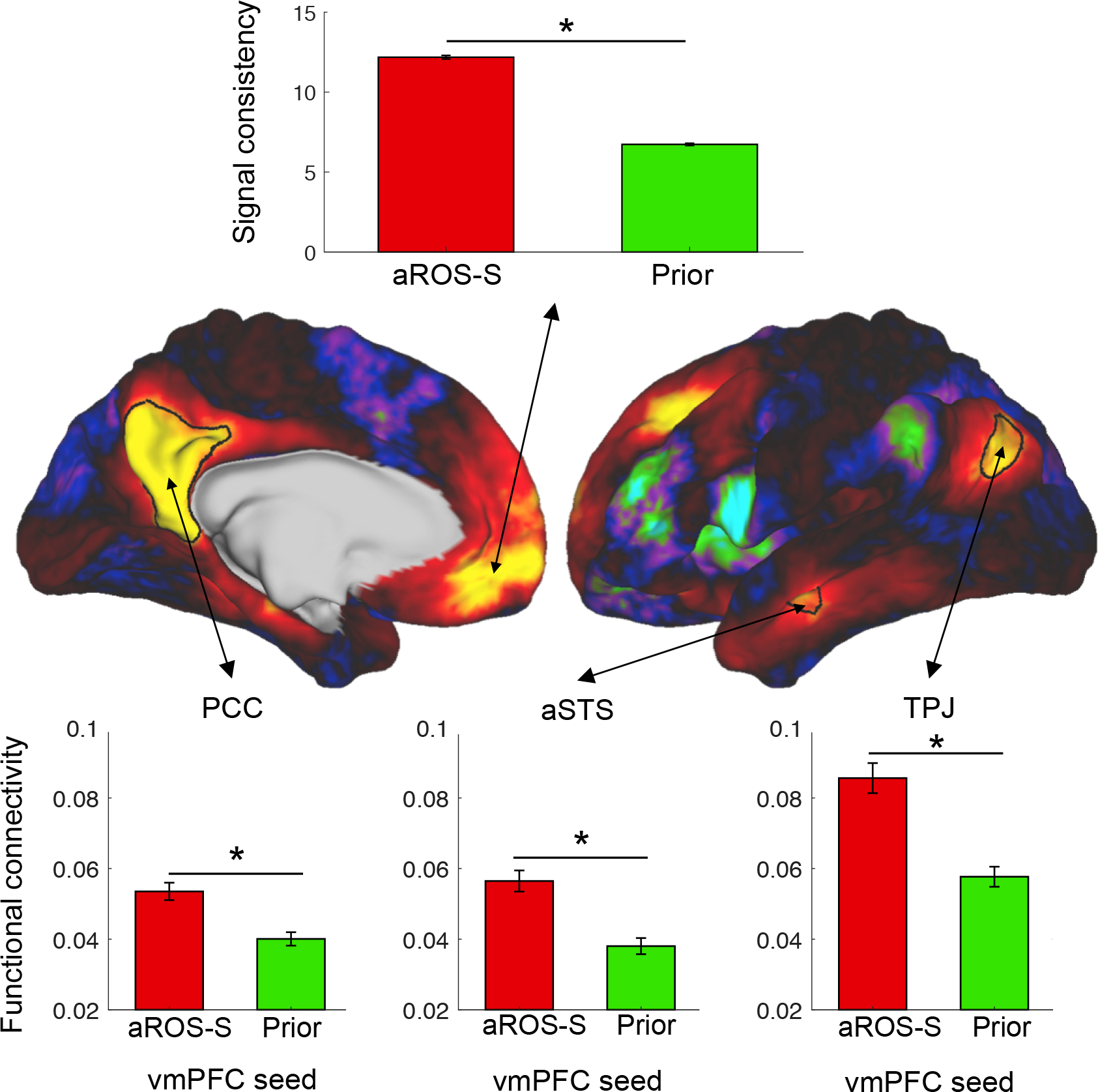
Default Mode Network components comparison between prior and anatomical ROI. **Top**. Signal consistency within aROS-S and ‘Prior’ ROI. Signal consistency is the inverse of the standard deviation of the raw signal across vertices averaged across time. **Middle**. Average Default Mode Network weights across left hemispheres (Right hemisphere not shown for illustrative purposes). The map was used to draw the PCC, aSTS and TPJ regions of interest. **Bottom**. Average functional connectivity across all hemispheres between the vmPFC seed (aROS-S or ‘Prior’) and the PCC (left), the aSTS (middle) and the TPJ (right). PCC: Posterior Cingulate Cortex; aSTS: anterior Superior Temporal Sulcus; TPJ: Temporo-Parietal Junction.

## DISCUSSION

The present study examined the sulcal morphology of the vmPFC and its relation to functional organization as reflected in resting state connectivity. We established that the superior rostral sulcus (ROS-S) and the supra-rostral sulcus (SU-ROS) can be found in all hemispheres and can thus be considered as the two principal sulci of the vmPFC. Organised around these two principal sulci of the vmPFC, we identified the probability of the presence of the supplementary medial sulci, such as the inferior rostral (ROS-I), the subgenual intralimbic (SILS) and the polar/supra-orbital sulci (MPS, VMPS, SOS, ASOS). We then demonstrated that the presence or absence of three supplementary sulci, namely the SILS, the ROS-I and PCGS had a major impact on the morphology of the vmPFC. First, the presence of a SILS induces an antero-ventral shift in the location of the SU-ROS. Second, the presence of ROS-I induces a dorsal shift of both the ROS-S and the SU-ROS. Thirdly, the SU-ROS merges with the PCGS, or with the CGS when the PCGS is not present, and this pattern has an impact on the location of the SU-ROS, although the shifts associated with the presence of SILS and ROS-I were greater than those induced by the presence of the PCGS. Moreover, the transformations applied to the brain volume images for registration to the MNI152 template and to surface-based maps for MSMAll registration (functional and sulcal surface registration) can be affected by the inter-individual variability in sulcal pattern. For example, we have shown that vmPFC sulci on the left hemisphere of brains registered to the MNI152 template do not correspond to the same sulci across subjects. This observation is of importance when considering interpreting the location of peaks of activity within the vmPFC.

The present results are consistent with those by Mackey and Petrides (2014) who examined cadaver brains on a smaller pool of subjects (n=13). In our study, the supra-rostral sulcus was observed in all hemispheres while they identified it in 91.3% of the hemispheres. This difference can be explained by a quite rare yet observed merging between the superior rostral sulcus and the supra-rostral sulcus. This merging can be observed either on the whole length of both sulci, resulting in a long and deep principal medial sulcus, or on a continuous merging between the antero-dorsal extremity of the ROS-S and the ventro-posterior extremity of the SU-ROS, resulting in an extremely long sulcus. Another noticeable difference is related to the SILS. We strictly used the SILS label only when no gyrus could be observed between the sulcus and the corpus callosum and we report a higher proportion of the presence of SILS than they do (44% against 30%). MPS, SOS and ASOS were observed in a high percentage of hemispheres, but we were able to identify an additional polar sulcus in 40% of the studied hemispheres. This sickle-shaped sulcus that occupies the most ventral part of the vmPFC is labelled as the ventral medial polar sulcus (VMPS). On average, its barycentre coordinates were y=58, z=−16. It is located just dorsal to the olfactory sulcus, and rostral to the ROS-I if a ROS-I is present.

Having determined the inter-individual variability in sulcal patterns and its impact on the vmPFC morphology, we then assessed whether sulcal morphology could influence the functional organization of the vmPFC and whether it could be used to improve characterization of functional activity observed in the vmPFC. For this purpose, we selected the Default Mode Network activation as a tool since the cluster of activity observed at the group level is covering a large part of the vmPFC. Importantly, we observed that the main hub of the Default Mode Network in the vmPFC is located in the anterior part of the ROS-S, regardless of the presence or absence of the secondary medial sulcus ROS-I. Note that the presence/absence of the ROS-I is variable across subjects and has a strong influence on the ROS-S location. In other words, the vmPFC peak of the functional DMN network is not optimally-defined by stereotaxic MNI coordinates, nor by any other volumetric or surface based group-averaged template. Instead, importantly, we have shown that the vmPFC peak of the Default Mode Network is best defined by *an anatomical sulcal feature* identified on an individual-by-individual basis.

The present results also raise methodological issues with regard to the investigation of the Default Mode Network, and to a greater extent, of the function of the vmPFC. Indeed, as in the interpretation of the location of activity in the MNI space, using a region of interest based on MNI coordinates or on vertices can influence the results. Sulcal features might be more accurate and our results call for the development of tools allowing an automatic detection of such features to characterize better functional subdivisions of the vmPFC.

Previous studies had shown that the location of functional activity in fMRI can often be predicted by the local morphology of the sulci in the frontal cortex (Amiez et al., 2006, 2013; Amiez and Petrides, 2014; Li et al., 2015; Amiez and Petrides, 2018). Morphological features not only explain cross-subject variability in functional connectivity but also explain behavioural measures (Bijsterbosch et al., 2018). However, to the best of our knowledge, this is the first time that it is shown that the functional organization of the vmPFC in resting state can be predicted by its local sulcal morphology. It should be noted that Mackey and Petrides (2014) have examined the sulcal patterns of the vmPFC in relation to cytoarchitecture and demonstrated that there is a good relationship between the sulci and particular cytoarchitectonic areas. Based on the location of the DMN peak in the rostral part of the ROS-S, it is therefore likely that the core node of the DMN in the medial prefrontal cortex is associated with a granular cortex area 14m or area 10m (Bludau et al., 2014; Mackey and Petrides, 2014). Supporting this anatomical correspondence between the vmPFC DMN node and area 10m is the fact that both have been functionally associated with supporting socio-cognitive functions (Mars et al., 2012). Anterior cingulate cortex (ACC) contributions to the DMN have also been reported in the literature (Vincent et al., 2007; Mantini et al., 2011; Qin and Northoff, 2011). However, our results would suggest that the implication of the ACC to the DMN is related to the strong monosynaptic connections of cingulate 24 and 32 with the frontopolar cortex (Petrides and Pandya, 2007). Future investigation should now be directed to extending this finding to other task-related functional activity in other subdivisions of the vmPFC, such as in tasks examining responses to decision variables (Lopez-Persem et al., 2016) or subjective valuation (Lebreton et al., 2009).

In conclusion, the present study provides a description of the variability of the vmPFC sulcal morphology across individuals and demonstrates that this variability might reduce the anatomical precision of group-average analyses. Thus, taking into account the local morphology of the vmPFC at the individual level to define regions of interest might be considered when assessing the multiple functions of this heterogeneous brain region.

## ACKNOWLEDGEMENT

Data were provided by the Human Connectome Project, WU-Minn Consortium (Principal Investigators: David Van Essen and Kamil Ugurbil; 1U54MH091657) funded by the 16 NIH Institutes and Centers that support the NIH Blueprint for Neuroscience Research; and by the McDonnell Center for Systems Neuroscience at Washington University. ALP received a fellowship from the Fondation pour la Recherche Medicale. JS was supported by a Sir Henry Dale Wellcome Trust Fellowship (105651/Z/14/Z). The Wellcome Centre for Integrative Neuroimaging is supported by core funding from the Wellcome Trust (203139/Z/16/Z).

## BIBLIOGRAPHY

Amiez C, Kostopoulos P, Champod A-S, Petrides M (2006) Local Morphology Predicts Functional Organization of the Dorsal Premotor Region in the Human Brain. J Neurosci 26:2724–2731.

Amiez C, Neveu R, Warrot D, Petrides M, Knoblauch K, Procyk E (2013) The Location of Feedback-Related Activity in the Midcingulate Cortex Is Predicted by Local Morphology. J Neurosci 33:2217–2228.

Amiez C, Petrides M (2014) Neuroimaging evidence of the anatomo-functional organization of the human cingulate motor areas. Cereb Cortex 24:563–578.

Amiez C, Petrides M (2018) Functional rostro-caudal gradient in the human posterior lateral frontal cortex. Brain Struct Funct 223:1487–1499.

Bartra O, McGuire JT, Kable JW (2013) The valuation system: A coordinate-based meta-analysis of BOLD fMRI experiments examining neural correlates of subjective value. NeuroImage 76:412–427.

Beckmann CF, Smith SM (2004) Probabilistic Independent Component Analysis for Functional Magnetic Resonance Imaging. IEEE Trans Med Imaging 23:137–152.

Bijsterbosch JD, Woolrich MW, Glasser MF, Robinson EC, Beckmann CF, Van Essen DC, Harrison SJ, Smith SM (2018) The relationship between spatial configuration and functional connectivity of brain regions. eLife 7.

Bludau S, Eickhoff SB, Mohlberg H, Caspers S, Laird AR, Fox PT, Schleicher A, Zilles K, Amunts K (2014) Cytoarchitecture, probability maps and functions of the human frontal pole. NeuroImage 93:260–275.

Bonnici HM, Chadwick MJ, Lutti A, Hassabis D, Weiskopf N, Maguire EA (2012) Detecting representations of recent and remote autobiographical memories in vmPFC and hippocampus. J Neurosci Off J Soc Neurosci 32:16982–16991.

Chiavaras MM, Petrides M (2000) Orbitofrontal sulci of the human and macaque monkey brain. J Comp Neurol 422:35–54.

Evans AC, Collins DL, Mills SR, Brown ED, Kelly RL, Peters TM (1993) 3D statistical neuroanatomical models from 305 MRI volumes. In: 1993 IEEE Conference Record Nuclear Science Symposium and Medical Imaging Conference, pp 1813–1817 vol.3.

Fischl B (2012) FreeSurfer. NeuroImage 62:774–781.

Fonov V, Evans A, McKinstry R, Almli C, Collins D (2009) Unbiased nonlinear average age-appropriate brain templates from birth to adulthood. NeuroImage 47:S102.

Friston KJ ed. (2007) Statistical parametric mapping: the analysis of funtional brain images, 1st ed. Amsterdam; Boston: Elsevier/Academic Press.

Glasser MF, Coalson TS, Robinson EC, Hacker CD, Harwell J, Yacoub E, Ugurbil K, Andersson J, Beckmann CF, Jenkinson M, Smith SM, Van Essen DC (2016) A multi-modal parcellation of human cerebral cortex. Nature 536:171–178.

Glasser MF, Sotiropoulos SN, Wilson JA, Coalson TS, Fischl B, Andersson JL, Xu J, Jbabdi S, Webster M, Polimeni JR, Van Essen DC, Jenkinson M (2013) The minimal preprocessing pipelines for the Human Connectome Project. NeuroImage 80:105–124.

Hänsel A, von Känel R (2008) The ventro-medial prefrontal cortex: a major link between the autonomic nervous system, regulation of emotion, and stress reactivity? Biopsychosoc Med 2:21.

Jenkinson M, Beckmann CF, Behrens TEJ, Woolrich MW, Smith SM (2012) FSL. NeuroImage 62:782–790.

Lebreton M, Jorge S, Michel V, Thirion B, Pessiglione M (2009) An Automatic Valuation System in the Human Brain: Evidence from Functional Neuroimaging. Neuron 64:431–439.

Li Y, Sescousse G, Amiez C, Dreher J-C (2015) Local Morphology Predicts Functional Organization of Experienced Value Signals in the Human Orbitofrontal Cortex. J Neurosci 35:1648–1658.

Lopez-Persem A, Domenech P, Pessiglione M (2016) How prior preferences determine decision-making frames and biases in the human brain. eLife 5:e20317.

Mackey S, Petrides M (2014) Architecture and morphology of the human ventromedial prefrontal cortex. Eur J Neurosci 40:2777–2796.

Mantini D, Gerits A, Nelissen K, Durand J-B, Joly O, Simone L, Sawamura H, Wardak C, Orban GA, Buckner RL, Vanduffel W (2011) Default Mode of Brain Function in Monkeys. J Neurosci 31:12954–12962.

Marcus D, Harwell J, Olsen T, Hodge M, Glasser M, Prior F, Jenkinson M, Laumann T, Curtiss S, Van Essen D (2011) Informatics and Data Mining Tools and Strategies for the Human Connectome Project. Front Neuroinformatics 5.

Marcus DS et al. (2013) Human Connectome Project informatics: Quality control, database services, and data visualization. NeuroImage 80:202–219.

Mars RB, Neubert F-X, Noonan MP, Sallet J, Toni I, Rushworth MFS (2012) On the relationship between the “default mode network” and the “social brain.” Front Hum Neurosci 6.

Noonan MP, Walton ME, Behrens TEJ, Sallet J, Buckley MJ, Rushworth MFS (2010) Separate value comparison and learning mechanisms in macaque medial and lateral orbitofrontal cortex. Proc Natl Acad Sci 107:20547–20552.

Passingham RE, Stephan KE, Kötter R (2002) The anatomical basis of functional localization in the cortex. Nat Rev Neurosci 3:606–616.

Petrides M (2002) The Mid-ventrolateral Prefrontal Cortex and Active Mnemonic Retrieval. Neurobiol Learn Mem 78:528–538.

Petrides M (2018) Atlas of the Morphology of the Human Cerebral Cortex on the Average MNI Brain, 1st Edition. Academic Press, new York.

Petrides M, Baddeley A (1996) Specialized Systems for the Processing of Mnemonic Information within the Primate Frontal Cortex [and Discussion]. Philos Trans Biol Sci 351:1455–1462.

Petrides M, Pandya DN (2007) Efferent Association Pathways from the Rostral Prefrontal Cortex in the Macaque Monkey. J Neurosci 27:11573–11586.

Qin P, Northoff G (2011) How is our self related to midline regions and the default-mode network? NeuroImage 57:1221–1233.

Raichle ME (2015) The Brain’s Default Mode Network. Annu Rev Neurosci 38:433–447.

Smith SM et al. (2013) Resting-state fMRI in the Human Connectome Project. NeuroImage 80:144–168.

Uğurbil K et al. (2013) Pushing spatial and temporal resolution for functional and diffusion MRI in the Human Connectome Project. NeuroImage 80:80–104.

Van Essen DC et al. (2012) The Human Connectome Project: a data acquisition perspective. NeuroImage 62:2222–2231.

Vincent JL, Patel GH, Fox MD, Snyder AZ, Baker JT, Van Essen DC, Zempel JM, Snyder LH, Corbetta M, Raichle ME (2007) Intrinsic functional architecture in the anaesthetized monkey brain. Nature 447:83–86.

Wallis JD (2011) Cross-species studies of orbitofrontal cortex and value-based decision-making. Nat Neurosci 15:13–19.

